# N-glycosylation Enables Smut Fungal Nge1 Orthologs to Prevent the Escape of Maize Evolved-PMEIs

**DOI:** 10.1101/2025.04.01.646729

**Authors:** Chibbhi Bhaskar, Minh-Quang Chau, Wei-Lun Tsai, Ooi-Kock Teh, Lay-Sun Ma

## Abstract

Cell wall integrity (CWI) is regulated by the coordinated activity of CW-modifying enzymes, including pectin methylesterases (PMEs) and their inhibitors (PMEIs). PMEs demethylesterify pectins, making them more susceptible to degradation and loosening the CW, facilitating pathogen invasion. Conversely, PMEIs inhibit PMEs, reinforcing the CWI and enhancing plant defense. However, how biotrophic pathogens overcome PMEI-mediated defense remains unclear. Here, we report that smut fungal effectors have evolved to directly target host specific PMEIs, manipulating cell wall integrity to enhance virulence. N-glycosylated Effector 1 (Nge1) from *Ustilago maydis* selectively interact with PMEI45 and PMEI46, as well as the auto-inhibitory PRO-domains of PMEs. This interaction disrupts PMEI inhibition, liberating PME19 and PME20, which reduce pectin methylesterification and likely loosen the CW, promoting fungal invasion. Notably, the interaction between Nge1 and PMEI45, but not PMEI46, is N-glycosylation-dependent. Restoring glycosylation in a non-glycosylated Nge1 ortholog allows it to functionally replace *U. maydis* Nge1, suggesting that smut fungal effectors have evolved through glycan modifications to overcome host-adapted PMEIs that would otherwise escape non-glycosylated effectors and impede fungal infection. Our findings reveal bidirectional host-pathogen strategies in a co-evolutionary arms race to fine-tune molecular interactions in the extracellular space.

## Introduction

The plant cell wall (CW) is a dynamic and complex structure that provides mechanical support and a physical barrier against pathogen invasion. Pectins, a major cell wall component, is a soluble and dynamic complex of polysaccharides rich in galacturonic acid ^1,2^. It forms a hydrated gel-like matrix in the space between cellulose microfibrils, determining the CW properties such as thickness, hydration, porosity, ion balance, and surface charge ^1,2^. The predominant form of pectin, homogalacturonan (HG), is synthesized with varying degrees of methylesterification, undergoes de-methylesterification by pectin methylesterases (PMEs) in the CW, to accommodate plant development and defense ^3^.

PMEs (E.C. 3.1.1.11), which belong to CAZy family 8 of carbohydrate esterases, are widespread in plants, bacteria, and fungi ^4,5^. In plants, there is a high diversity of PMEs, with counts ranging from 66 in *Arabidopsis thaliana* ^6^ and 43 in *Zea mays* ^7^ to 105 in *Linum usitatissimum* ^8^. They are classified into two types. Type 1 PME harbors an N-terminal auto-inhibitory domain (PRO) in addition to the PME enzymatic domain. Type 2 PME possesses only the enzymatic domain, similar to those found in bacteria and fungi^9^. PME modulates the degree of pectin methylesterification through two distinct mechanisms to influence CW flexibility: random and ordered block-wise de-methylesterification ^10^. Removal of methyl esters groups in a random manner will render the HG to become vulnerable to plant-or pathogen-derived pectic enzymes like polygalacturonases, which cause CW loosening. Alternatively, PME can remove methyl esters in an ordered block-wise manner and results in continuous stretches of de-methylesterified GalAs that favor Ca2+ binding to stiffen the CW.

Pathogens exploit the plant PMEs to facilitate their cellular penetration and proliferation. The expression of PMEs is reported to be upregulated during infection by pathogens such as *Fusarium oxysporum*, *Taphrina deformans*, *Botrytis cinerea*, and *Pectobacterium carotovorum* ^11–13^. Notably, *Ustilago maydis* significantly increases PME activity in susceptible maize lines more than that in resistant ones to influence the extent of pectin methylesterification during biotrophy ^14^. The distinct mechanisms of PME in modulating the structural dynamics of plant CWs highlight their intricate roles and importance during pathogen infection.

On the other hand, PME activity is modulated by PME inhibitors (PMEIs). Both the PMEIs and PRO-domains of PME display a network of α-helices and possess structural and functionality similarities in their ability to inhibit PMEs ^3^. It has been suggested that PMEIs may have derived from the duplication and divergence of PRO-domains and have undergone rapid evolution ^15^. Like PMEs, PMEIs are abundant in plants, with 71 homologs in *A. thaliana*, 49 in *Z. mays,* and *O. sativa* ^7,15,16^. Overexpression of PMEIs elevates the degree of pectin methylesterification and enhances plant resistance, thereby limiting the pathogen access, including *B. cinerea*, *Alternaria brassicicola*, *F. oxysporum*, *Pseudomonas syringae*, *Bipolaris sorokiniana*, *F. graminearum*, and *Verticillium dahlia* ^17–21^. However, the underlying mechanisms by which pathogens overcome the evolved PMEI’s defense during successful infection remain elusive.

*U. maydis* is a biotrophic smut fungus causing corn smut in maize. The pathogenicity is characterized by induction of anthocyanin and tumor formation in all aerial parts of plants. It secretes a large arsenal of effector proteins to facilitate infection. These effector proteins either localize in the apoplast or translocate into plant cells to suppress plant immunity and reprogram plant physiology, ultimately establishing an intimate relationship with the host ^22,23^. However, only a few effectors have been functionally characterized. Given that *U. maydis* increases PME activity during infection ^14^, *U. maydis* may employ effectors to alter plant CW architecture by modulating the PME activity to facilitate its penetration.

In this study, we demonstrate that *U. maydis* manipulates the degree of pectin methylesterification in host by interfering with the activity of PMEIs. This is accomplished by deploying a secreted N-glycosylated Effector 1 (Nge1) that targets multiple PMEIs in the host to alleviate PMEs from inhibition. This mechanism results in a pronounced impact on the pectin methylesterification, potentially influencing the cell wall flexibility. Our study has uncovered a unique strategy employed by the smut fungal pathogens in which glycan decoration on a novel effector could alter the dynamics of host cell wall instead of degradation, adding a layer of sophistication to their pathogenicity strategy.

## Results

### Nge1 is an apoplast N-glycosylated virulence factor

*Ustilago maydis* harbors 53 novel core effectors within its genome, including N-glycosylated effector 1 (Nge1), a singleton encoded by *UMAG_11362,* which contributes to the virulence ^24^. Nge1 is a 113 amino acids-long peptide and features an N-terminal signal peptide (SP), a putative N-glycosylation site located at the 71^st^ asparagine (N) position (denoted as N71), followed by an intrinsically disordered region (IDR) (Figure 1A). This protein lacks any distinct functional domains as predicted by InterPro ^25^. The structure predicted by AlphaFold exhibits no resemblance to any proteins in the AlphaFold protein structure database (Figure S1A). qRT-PCR analyses revealed that *NGE1* was not expressed in axenic culture but its expression was elevated during the biotrophic stages of *U. maydis* SG200, a solopathogenic haploid strain ^26^ (Figure 1B). Deletion of *NGE1* in SG200 led to a notable decrease in disease symptoms in infected maize seedlings (Figure 1C), which was complemented by expressing a single copy of C-terminal HA-tagged *NGE1* under the control of its native promoter. However, abolishing the N-glycosylation site by an asparagine to glutamine substitution at the N71 site (N71Q) did not restore the virulence phenotype of the Δ*nge1* mutant. These findings indicate that Nge1 contributes to the virulence of *U. maydis,* likely in an N-glycosylation-dependent manner and prompted further investigation.

**Figure 1.**
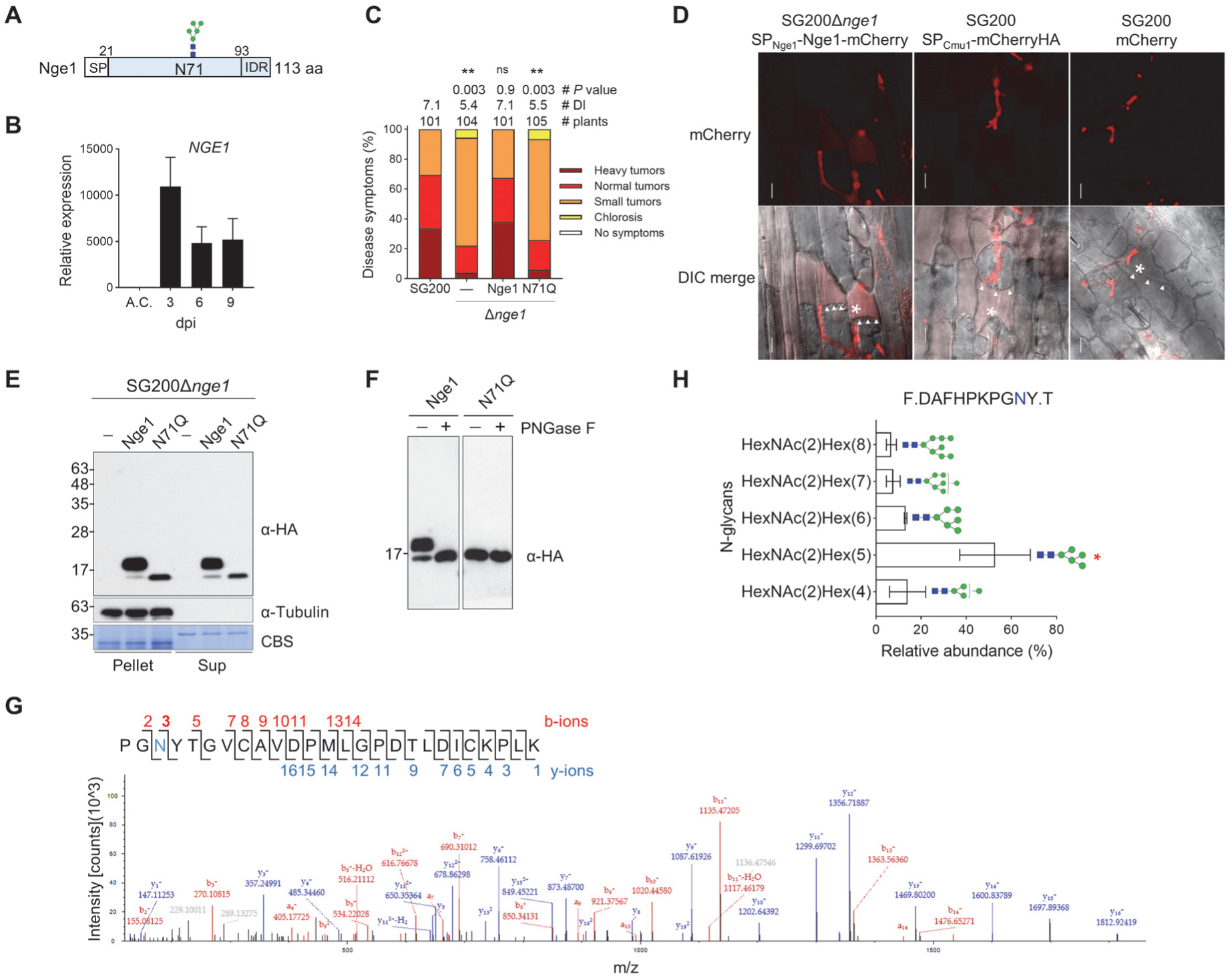
Nge1 is an apoplast N-glycosylated virulent factor. (**A**) Schematic drawing of *U. maydis* Nge1 (UMAG_11362) protein. SP: signal peptide. IDR: intrinsic disordered region. The putative N-glycosylated asparagine N71 is indicated with the predominant N-glycan form identified. (**B**) *NGE1* expression during biotrophy. Total RNA samples prepared from SG200 infected leaves at indicated time points and from cells cultured in liquid medium (A.C) were subjected to qRT-PCR analysis. Expression levels of *NGE1* were normalized relative to the peptidyl-prolyl isomerase (*PPI*) gene. Values are mean ± standard deviation (SD) of three biological replicates. (**C**) Virulence of the indicated strains was assessed. Complemented strains expressing C-terminal HA-tagged Nge1 and glycosylation mutant N71Q were driven by *Nge1* native promoter. Maize seedlings were infected with the indicated strains and disease symptoms were scored at 8-day post infection (dpi) following the disease scoring scheme developed^26^. The average disease index (DI) of three biological replicates are shown. Significant differences compared to SG200 were determined by a two-tailed unpaired Student’s *t*-test (\**P* < 0.05; \*\**P* < 0.01; \*\*\**P* < 0.001). ns: no significance. (**D**) Nge1 localizes to the apoplast. Maize leaves infected by strains expressing indicated proteins with or without SP were plasmolyzed using 0.8 M mannitol. SP_Cmu1_: SP derived from *U. maydis* chorismate mutase Cmu1 ^32^. Asterisks and triangles indicate the enlarged apoplastic space and shrunken plasma membrane respectively. Bars 10μm. (**E**) Nge1 is a glycosylated secreted protein. Proteins from cell pellet and supernatant fractions of SG200Δ*nge1* and SG200Δ*nge1* constitutively expressing Nge1-HA and N71Q-HA, driven by *otef* promoter, were subjected to immunoblotting. Tubulin served as a non-secreted control and Coomassie blue staining (CBS)-membranes were used to verify loading. A representative result from three independent experiments is shown. (**F**) Deglycosylation of Nge1. Supernatant fractions of *U. maydis* constitutively expressing Nge1-HA and N71Q-HA were treated with or without PNGase F and subjected to immunoblotting. A representative result from at least two independent experiments is shown. (**G**) Identification of N-glycosylation site N71 by LC-MS/MS. The detected b and y-ions of MS/MS fragmentation of the peptide “PG**N**YTGVCAVDPMLGPDTLDICKPLK are annotated. The b-ions fragments (b3 to b14), showing an increase of +0.984 Da due to the deamidated asparagine (N) residue post PNGase F treatment, are indicated in bold. (**H**) Detection of N-glycans by LC-MS/MS. The graph illustrates the average relative abundance percentage of detected N-glycan forms on the glycopeptide FDAFHPKPG**N**YT across three biological replicates. The column marked with an asterisk represents the predominant form identified. HexNAc and Hex is proposed as GlcNAc and mannose respectively. Blue square, *N*-acetylglucosamine; Green circle, mannose. The number of sugar molecules is indicated.

Nge1 is predicted to localize to the extracellular space by the ApoplastP ^27^. To test this, we examined Nge1 localization in the Δ*nge1*-Nge1-mCherry infected leaves. Fluorescence microscopy revealed that Nge1-mCherry localized in the enlarged apoplastic space caused by plasmolysis (Figure 1D). This localization pattern was consistent with that of the positive control, SP-mCherry, whereas the cytosolic mCherry remained intracellular in the fungal cells. Immunoblot analysis of apoplast fluid from maize leaves infected with Δ*nge1_*Nge1-mCherry also detected full-length fusion proteins (Figure S1B), confirming Nge1 secretion and apoplastic localization.

To analyze the migration pattern of proteins, we constitutively expressed C-terminal HA-tagged Nge1 and N71Q in Δ*nge1*, under the control of the *otef* promoter in axenic culture (Figure 1E). In Δ*nge1_*Nge1, a predominant band migrating above 17 kDa and a minor band below 17 kDa were detected in both the cell pellet and supernatant fractions. In contrast, only a single band below 17 kDa was observed in the Δ*nge1_*N71Q fractions. Peptide-*N*-glycosidase F (PNGase F) digestion Nge1, which specifically removes N-glycans, yielded a single band with a migration pattern akin to that of N71Q (Figure 1F). These findings indicate that the protein band migrating above 17 kDa is N-glycosylated.

To confirm the N-glycosylation site, we conducted Liquid Chromatography-Tandem Mass Spectrometry (LC-MS/MS) analysis on PNGase F-treated Nge1-mCherryHA (Figure S1C). PNGase F treatment induces deamination of asparagine, resulting in a mass increase of 0.985 Da, which can be detected by LC-MS/MS analysis^28^. The fragmented b3+ ion of the detected peptide, PG**N**YTGVCAVDPMLGPDTLDICKPLK, showed a mass-to-charge ratio of (m/z) of 270.108, which is 0.985 Da higher than the theoretical mass value of 269.123 m/z, clearly indicating a mass shift at position N71 (Figure 1G). This shift was consistently observed in the subsequent b-ion fragments, confirming N-glycosylation at N71 in Nge1.

To investigate the N-glycan pattern, affinity-purified Nge1 proteins were treated with chymotrypsin, which cleaves at aromatic amino acids to yield the glycopeptide FDAFHPKPG**N**YT containing the N71 glycosite, followed by LC-MS/MS analysis ^29^ (Figure 1H and Figure S1D). Our results revealed that the glycopeptide possessed oligomannose-type N-glycan structures with varying numbers of mannose units at the N71 position, among which HexNAc(2)Hex(5) was the predominant N-glycan form identified. These findings collectively support the notion that the N71 residue of Nge1 is modified with oligomannose-type N-glycans.

### Nge1 targets PMEI45 and its homologs

Having shown that Nge1 localized at the extracellular space, we next sought to identify its target within this space. We purified Nge1 or Nge1(N71Q) from culture supernatants of *U. maydis* strains and incubated them with apoplastic fluid from Δ*nge1*-infected maize leaves to capture respective interactors. Distinct protein bands were detected in the Nge1 sample, but was absent in the Nge1(N71Q). The sample bands were excised, combined, and subjected to LC-MS/MS analysis (Figure S2A). Four candidates associated with pectin methylesterification regulation but not in the Nge1(N71Q) sample were exclusively detected in the Nge1 sample (Figure S2B). These include a type-I PRO-domain-containing PME19 (PRO-PME19; B8A2X5) ^7^, PMEI45 (A0A1D6HA45) ^7^, and two cell-wall DUF642 domain-containing proteins (B4FRA6 and A0A1D6H3D4) ^30^. We further validated Nge1 interaction with these candidates using yeast-two hybrid (Y2H) analysis and confirmed that Nge1 interacted with both PMEI45 and PRO19 (the PRO-domain of PME19; Figure 2A), but not with the full-length PRO-PME19 and DUF642 proteins (Figure S2C-E). These results demonstrated that Nge1 targets the PME inhibitors, PRO19 and PMEI45.

**Figure 2.**
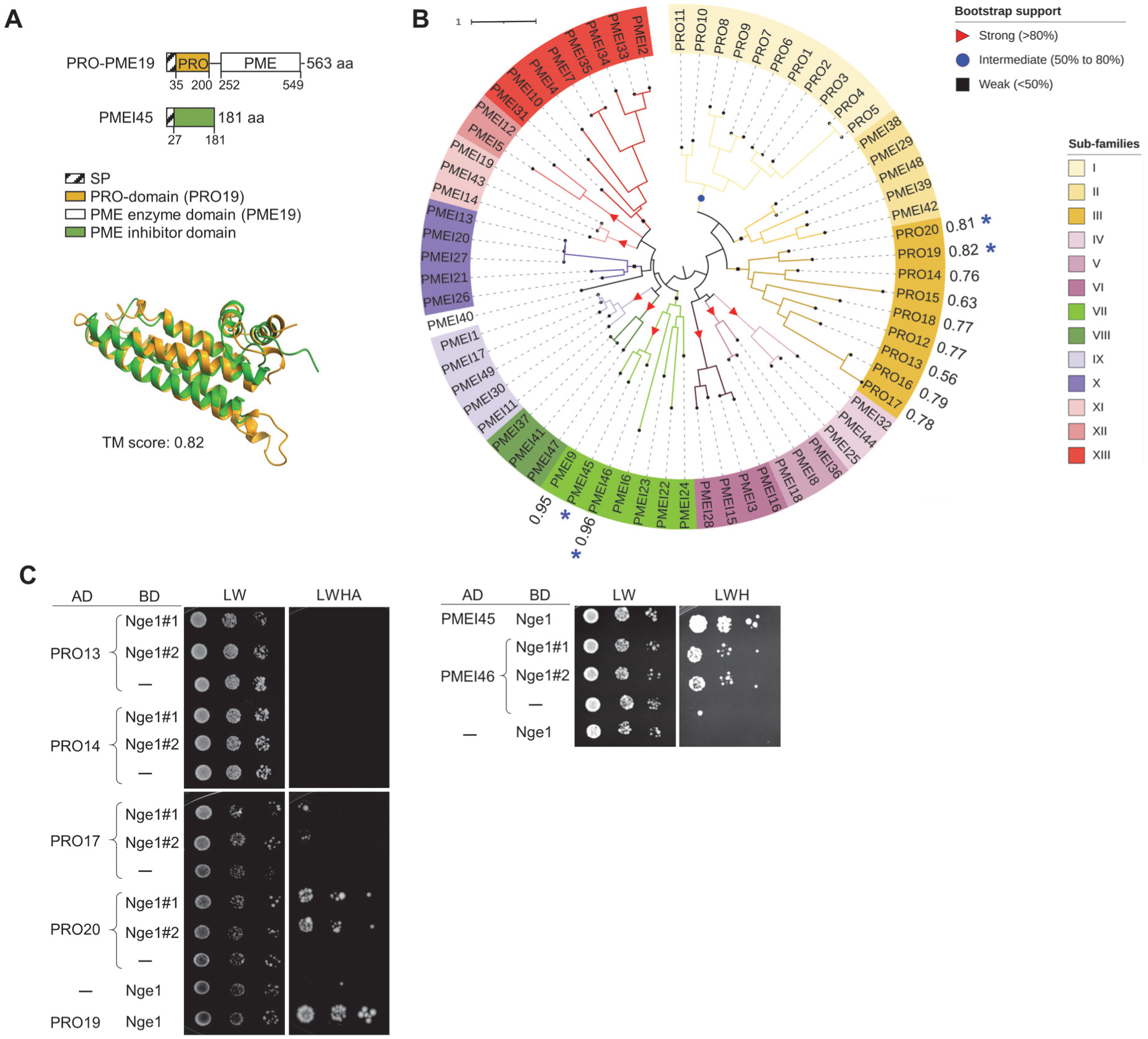
Nge1 targets PMEI45 and its homologs. (**A**) PMEI45 and PRO-domain of PME19 exhibit structural similarity. Domain architecture and superimposition of maize PRO19 and PMEI45. The position of PRO and PMEI is indicated. Superimposition of AlphaFold structures of PMEI45 (green) and PRO19 (gold) domains, excluding signal peptide (SP), was performed using ChimeraX, with the TM-align score indicated. (**B**) Phylogeny of PMEI domains of 49 PMEIs and PRO-domains of type-I PMEs, annotated by UniProt. Amino acid sequences aligned using Clustal Omega and the phylogenetic tree was constructed using PhyML 3.0 with the Q. pfam model and 1000 bootstraps. The strength of the bootstrap values is indicated at nodes using symbols (red triangle; >80%, blue circle; 50-80% and black square; <50%). The proteins are separated into 13 sub-families. TM-scores from Figure S3B-C, are indicated next to the protein superimposed with PMEI45. Accession # of proteins is available in Supplementary Table S4. The clades carrying PMEI45 and PRO-19 are highlighted in green and gold respectively. Blue asterisks mark the PME inhibitors interacted with Nge1 determined by Y2H assays. (**C**) Nge1 interacts with PRO19, PRO20, PMEI45, and PMEI46. Yeast transformants containing two plasmids expressing indicated proteins fused to GAL4 activation domain (AD) or binding domain (BD) without signal peptide were grown on SD-Leu/-Trp (LW) or SD-Leu/-Trp/-His/-Ade (LWHA) plates for 2-3 days. ̶, empty vector; Cmu1 served as the positive control. Similar results were observed in at least two independent repeats. Detailed information on the protein regions included in the Y2H analysis is available in Table S5.

Since PMEIs may originate from the duplication and divergence of PRO-domains, we superimposed the AlphaFold structures of PMEI45 and the PRO19 domain to further investigate their structural folding. This analysis revealed a high degree of similarity, with a TM-align score of 0.82 ^31^ (Figure 2A). Given that the maize genome encodes a total of 20 type-I PMEs and 49 PMEIs ^7^, we then explored whether Nge1 can target homologs with high structural similarity to PMEI45. We combined phylogenetic tree analysis with AlphaFold pairwise alignment to investigate this possibility. In the PMEI phylogenetic tree, all PRO-domains of type-I PMEs and the 49 PMEIs are clustered into 13 subfamilies, with PRO19 and PMEI45 located in subfamilies III and VII, respectively (Figure 2B). Based on the expression profile compiled from RNA-seq data of *U. maydis*-infected samples ^22^, *PMEI45* was among the upregulated genes during infection (Figure S3A). Notably, *PME19* exhibited the highest induction among subfamily III members, suggesting that *U. maydis* may upregulate this gene to facilitate its infection.

To assess whether members of these two subfamilies could be additional targets of Nge1, we used PMEI45 as bait and superimposed its AlphaFold structures with all PRO domains and PMEIs, revealing varying degrees of structural similarities. TM-align scores ranged from 0.56 to 0.81 for PRO-domains and up to 0.96 for PMEIs (Figures 2B and S3B-C). Among the candidates selected for Y2H screening, only PRO20 and PMEI46 interacted with Nge1 (Figure 2C). By integrating structure-alignment and Y2H analysis, we revealed that Nge1 can target multiple PME inhibitors, including PRO19, PRO20, PMEI45, and PMEI46.

### Glycosylated Nge1 specifically interacts with PMEI45

To verify these interactions, we used bimolecular fluorescence complementation (BiFC) approach by expressing split yellow fluorescent protein (YFP)-fused proteins for Nge1 and the PME inhibitors in *Nicotiana benthamiana* cells. YFP signals were clearly visible in leaves co-expressing PMEI45 and Nge1, whereas weak or undetectable signals were observed when PMEI45 was co-expressed with the chorismate mutase effector Cmu1 ^32^, which possesses similar oligomannose-type N-glycans (Figures 3A and S4A-B), illustrating that glycan binding is not the underlying factor for YFP reconstitution. Notably, despite similar protein expression levels between these samples (Figure S4C), Nge1(N71Q) expression resulted in a marked reduction in YFP intensity. This finding is consistent with the Y2H data (Figure S2D), highlighting the necessity of N-glycosylation for the Nge1-PMEI45 interaction.

**Figure 3.**
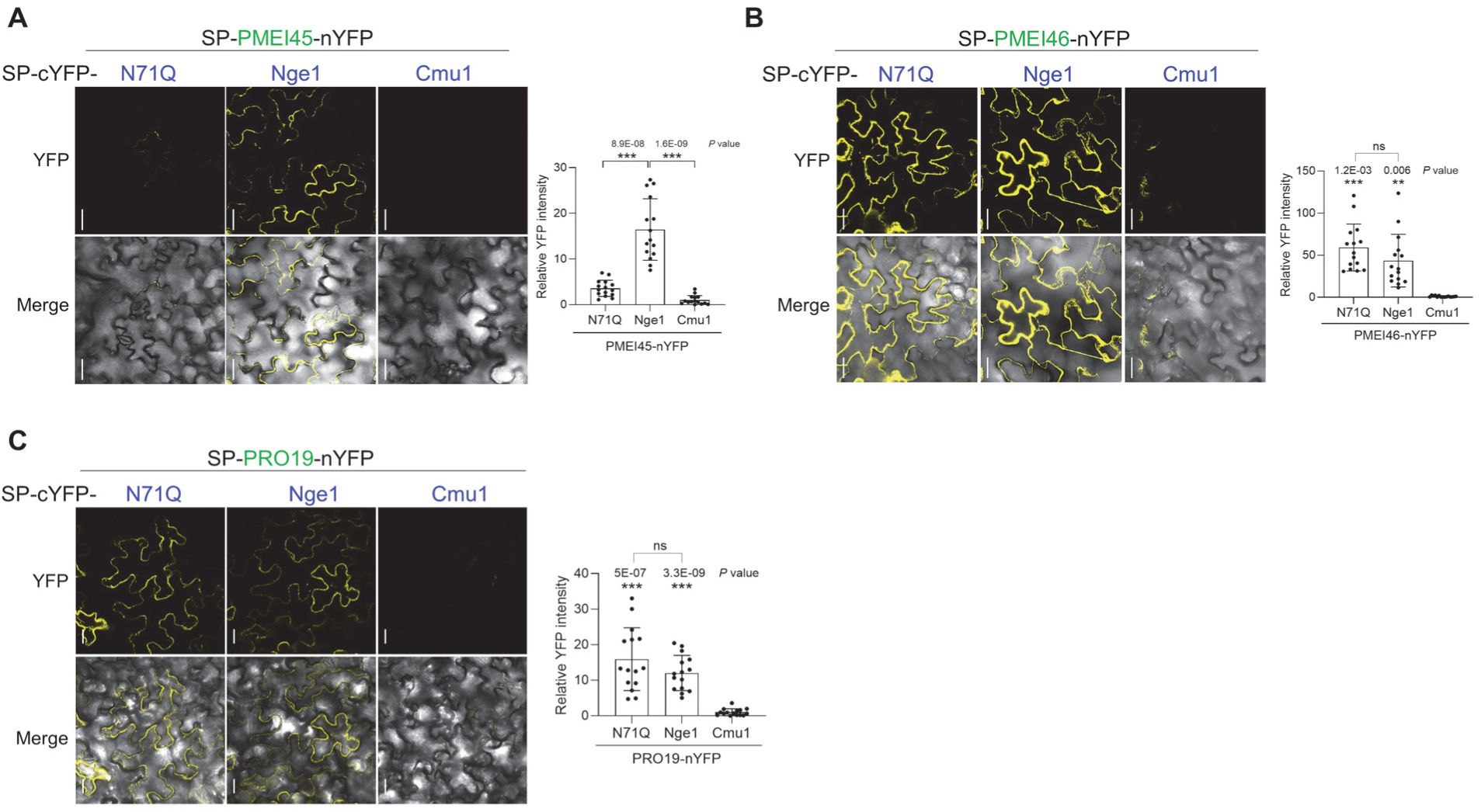
Nge1 displays a glycosylation-dependent interaction with PMEI45. BiFC analysis of Nge1 and N71Q interaction with PMEI45 (A), PMEI46 (B), and PRO19 (C) *in planta*. cYFP-Cmu1 served as a negative control. The SP of maize antifungal protein (AFP1) was used to deliver the indicated proteins to the apoplast ^54^. YFP epifluorescence image (514 nm) and that merged with differential interference contrast (DIC) image are shown. Fluorescent signals were quantified by ZEISS ZEN microscopy software. YFP fluorescence was normalized to cell wall autofluorescence. The graph depicted the fold differences relative to the negative control expressing indicated proteins and Cmu1. Values indicate mean ± SD from total samples obtained from three independent biological assays and dots depict the values of individual samples across these replicates. Significant differences compared to the control or between the samples were determined by a two-tailed unpaired Student’s *t*-test (\**P* < 0.05; \*\**P* < 0.01; \*\*\**P* < 0.001). All bars 50 μm.

Although Nge1 also interacted with PRO19 and PMEI46, their interaction was glycosylation-independent, as evidenced by the BiFC and Y2H assays (Figure 3B-C; Figure S2E & S4D-E). We therefore conclude that Nge1 interacts with these PME inhibitors, but displays specificity in glycosylation-dependent binding with PMEI45.

### Nge1 relieves PME19 from PMEI45-imposed inhibition

The high structural similarities among the identified PME inhibitors further suggest that PMEI45 and PMEI46 likely interact with PME19 and PME20 to exert inhibition. Consistently, PMEI45 and PMEI46 showed specific interactions with both PME19 and PME20, but not with the full-length PRO-PME19 and PME13 (Figure 4A-B and S5A-B). These findings align well with the AlphaFold pairwise structure alignment and suggest that PMEI45 and PMEI46 could functionally substitute for PRO19 and PRO20 in inhibiting PME19 and PME20 activity. Based on these findings, we propose that Nge1 could relieve at least PME19 and PME20 from the inhibition by PMEI45 and PMEI46 in an N-glycosylation-dependent and-independent manner, respectively.

**Figure 4.**
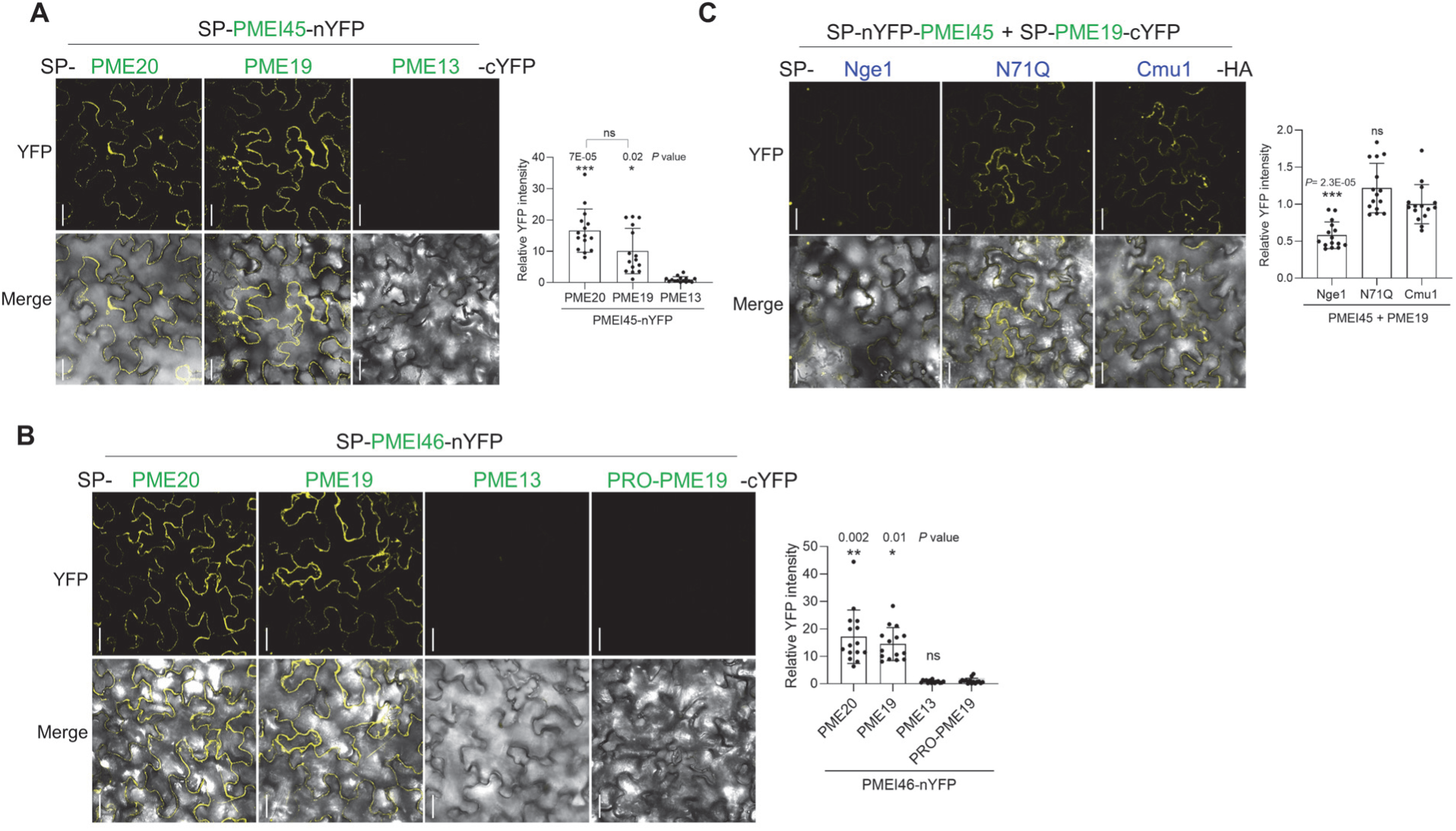
Nge1 alleviates PME19 inhibition by PMEI45 in a glycosylation-dependent manner. PMEI45 (**A**) and PMEI46 (**B**) interact with the PME19 and PME20 enzymatic domains. BiFC analysis of PMEI45 and PMEI46 interactions with the full-length PRO-PME19 and the enzymatic domains of indicated PMEs *in planta*. The graph depicted the fold differences relative to the control expressing PME13 or PRO-PME19. (**C**) Nge1 disrupts the PMEI45-PME19 interaction depending on glycosylation. BiFC analysis of PMEI45 interaction with PME19 in the presence of Nge1 and N71Q. Cmu1 served as a negative control. The YFP fluorescence was normalized to cell wall autofluorescence. The graph depicted the fold differences relative to the Cmu1 sample. Values indicate mean ± SD from total samples obtained from three independent biological assays and dots depict the values of individual samples across these replicates. Significant differences compared to Cmu1 as determined by a two-tailed unpaired Student’s *t*-test. All bars 50 μm.

To explore the impact of Nge1 on the interaction between PMEI45 and PME19, we co-expressed Nge1 with nYFP-PMEI45 and PME19-cYFP in BiFC assays. In contrast to the glycosylation-defective N71Q and Cmu1, the presence of Nge1 significantly reduced the YFP reconstitution, suggesting that PMEI45-PME19 interaction was compromised or weakened (Figure 4C and S5C). Importantly, the interfering effect of Nge1 is N-glycosylation-dependent. Our finding, therefore, support a scenario in which Nge1 interferes with the PMEI45-PME19 interaction and may consequently relieve PME19 from PMEI45-imposed inhibition to enhance pectin demethylesterification.

### Nge1 decreases methylesterified pectin levels to enhance virulence

Having shown that Nge1 could alleviate more than one PME from PMEI inhibition, we then investigated the impact of this rescue effect on the degree of pectin methylesterification ^14^. We assessed the degree of pectin methylesterification in cross-sections of maize leaves that expressed SP-Nge1-HA, as introduced by the means of Foxtail mosaic virus-mediated overexpression system (VOX) ^33^. Immunostaining with LM19 and JIM7 antibodies, which specifically detect low or high methylesterified pectin respectively, revealed that Nge1 increased the amount of pectin with low methylesterification at the cell wall compared to the negative control lines of SP-GFP (Figure 5A). Consistently, highly methylesterified pectins was greatly reduced in the Nge1-expressing line, as detected by the JIM7 antibody (Figure 5B).

**Figure 5.**
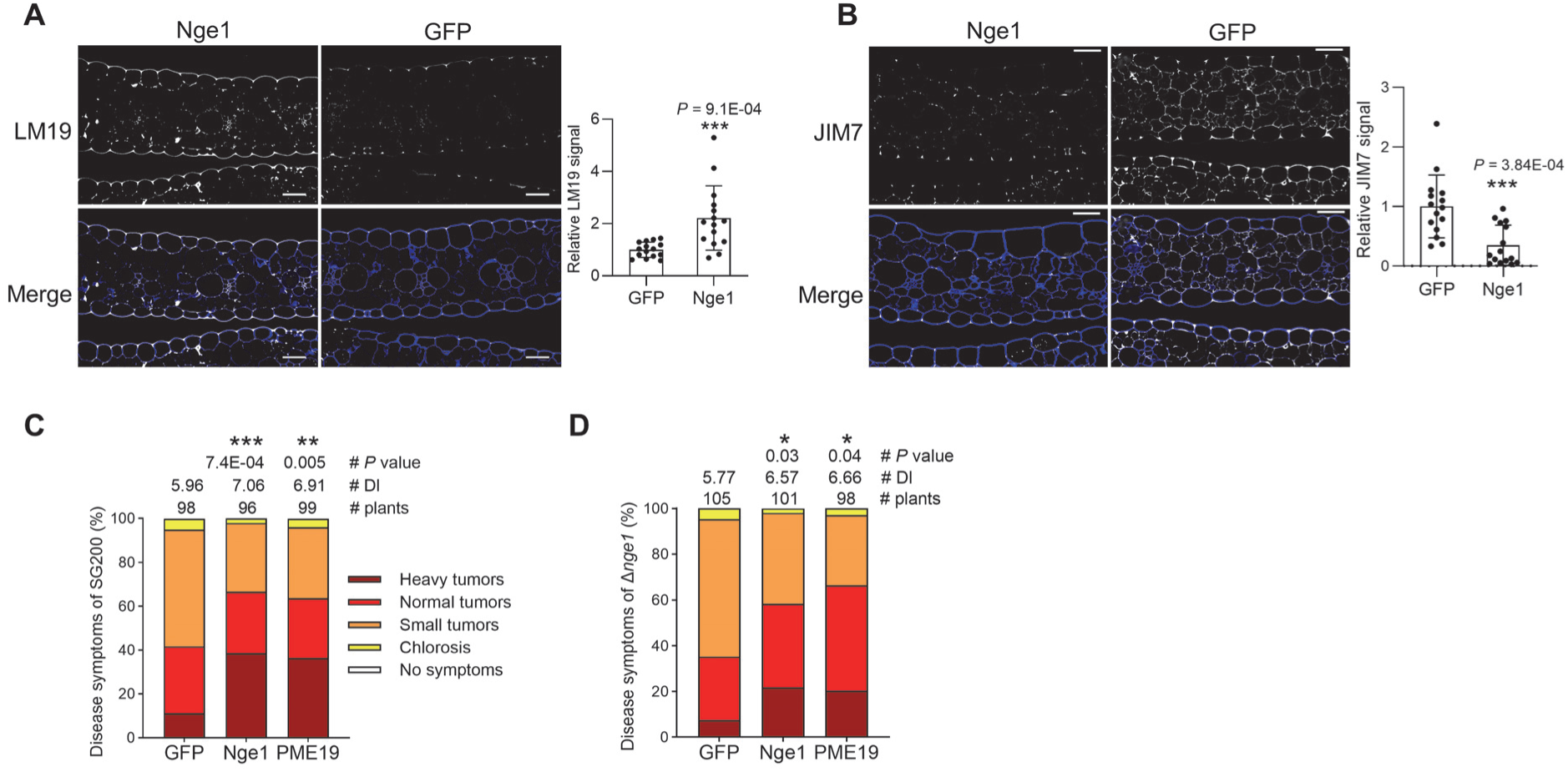
Nge1 increases the levels of low-methylesterified pectin to enhance virulence. (**A, B**) Immunostaining of pectins in cross-sections of maize leaves constitutively expressing SP_AFP1_-Nge1-HA or SP_AFP1_-GFP. Cell walls were stained with Calcofluor White. To detect pectins, rat monoclonal antibodies LM19 and JIM7 were used to detect de-esterified and methyl-esterified pectins, respectively, followed by immunostaining with a goat anti-rat secondary antibody conjugated to Alexa-Fluor 488. Fluorescent signals were visualized and quantified by ZEISS ZEN microscopy software. LM19 and JIM7 fluorescence was normalized to that of Calcofluor. The graph depicted the fold differences relative to the negative control, GFP. Biological replicates were conducted in triplicate, with each dot representing the value of an individual sample in the replicates. Significant difference compared to the control was determined by a two-tailed unpaired Student’s *t*-test (\**P* < 0.05; \*\**P* < 0.01; \*\*\**P* < 0.001). (**C, D**) Virulence of SG200 and Δ*nge1* on the VOX lines constitutively expressing SP_AFP1_-GFP, SP_AFP1_-Nge1-HA, SP_AFP1_-PME19-His. The number of infected plants and disease index (DI) from three biological replicates are shown.

To understand the biological relevance of modulating methylesterified pectin levels, we assessed the virulence levels of SG200 and Δ*nge1* in the SP-Nge1-HA, SP-PME19-His, and SP-GFP expressing maize lines (Figure 5C-D). In both the Nge1 or PME19 lines, the virulence of SG200 and Δ*nge1* was significantly enhanced compared to GFP lines. Collectively, our in vivo results suggest that Nge1 confers virulence by enhancing host susceptibility, likely through relieving the PMEs from PMEI to reduce methylated pectin in the wall, ultimately loosening the cell wall to facilitate penetration.

### Conserved N-glycosylation-dependent virulence function of Nge1 in smut fungi

Based on the glycosylation-dependent interaction between Nge1 and PMEI45, along with the phylogenetic divergence of PMEI45 from PMEI46, we hypothesize that PMEI45 may have evolved to evade recognition by non-glycosylated Nge1. This would ensure effective control over PME19 and PME20 activity, preventing the PMEs from being exploited by pathogens. We thus explored the conservation of Nge1 glycosylation status across smut fungi by conducting a BLASTp analysis ^34^. Amino acid sequence alignments of thirteen Nge1 orthologs identified across various fungal species highlighted conserved residues, showing a high degree of identity ranging from 44.34% to 62.92% (Figure 6A). The N-glycosylation site (NYT) of *U. maydis* UmNge1 is well-conserved in the orthologs of *Sporisorium reilianum f. sp. reilianum* (SrsNge1), *S. reilianum*_SRZ2 (SrzNge1), and *S. scitamineum* (SsNge1).

**Figure 6.**
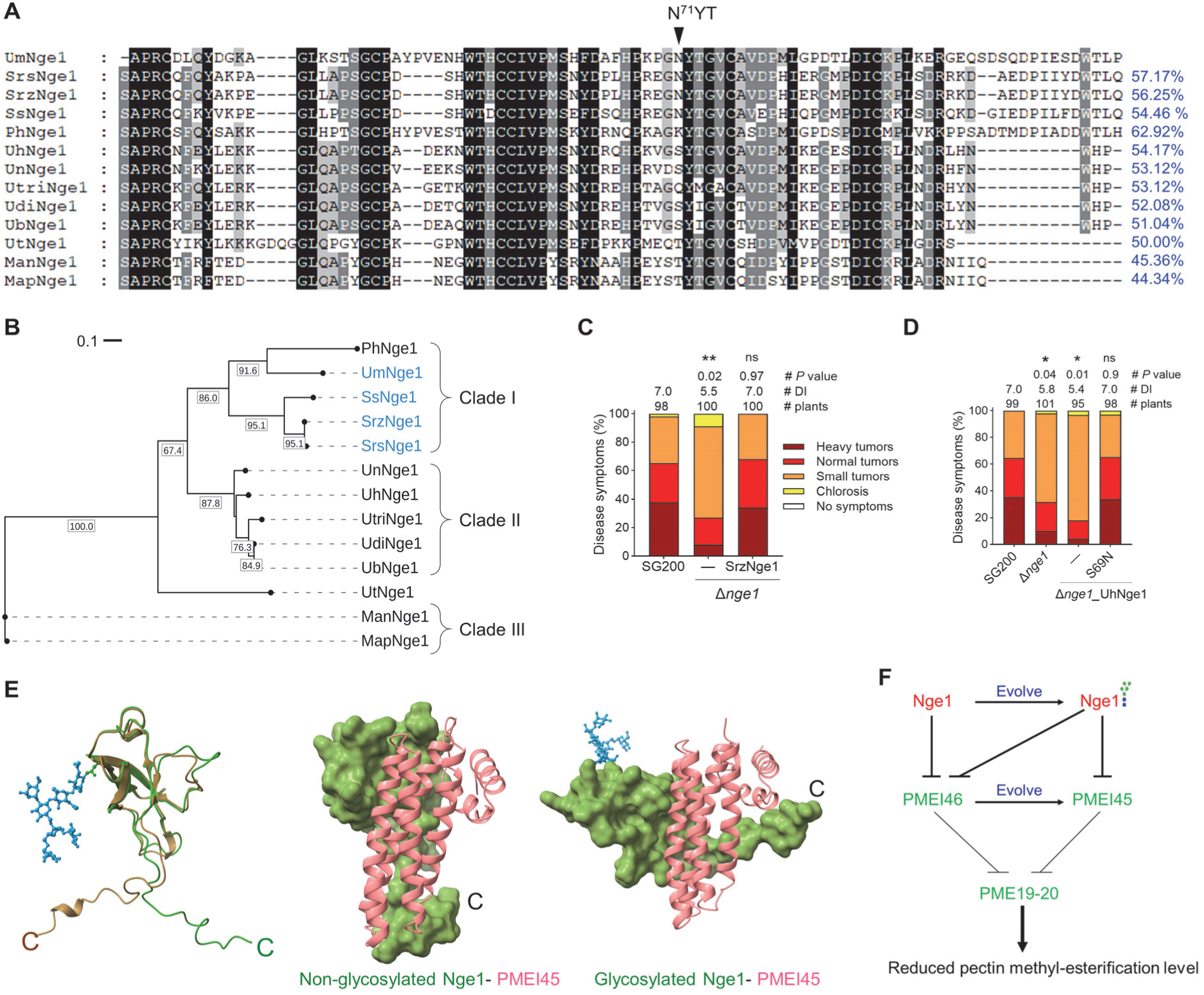
Nge1 exhibits conserved N-glycosylation-dependent virulence function in smut fungi. (**A**) Sequence alignment of Nge1 orthologs. Ortholog proteins with identity percentages to UmNge1 were identified through NCBI BLASTp analysis of the Non-redundant protein sequences database (https://blast.ncbi.nlm.nih.gov/Blast.cgi). Protein sequences without SP were aligned using Clustal Omega (https://www.ebi.ac.uk/jdispatcher/msa/clustalo), and visualized using GeneDoc. *U. maydis* (Um; XP_011388023.1) ^26^; *Sporisorium reilianum* f. sp*. reilianum* (Srs; SJX61796.1); *Sporisorium reilianum* f. sp*. zeae* (Srz; CBQ71923.1) ^66^; *Sporisorium scitamineum* (Ss; CDR99801.1) ^67^; *Pseudozyma hubeiensis* (Ph; XP_012189844.1) ^68^; *Ustilago hordei* (Uh; XP_041411439.1) ^69^; *Ustilago nuda* (Un; KAJ1027351.1) ^70^; *Ustilago tritici* (Utri; KAJ1029609.1) ^71^; *Ustilago diandrus* (Udi; SOV04500.1) ^72^; *Ustilago bromivora* (Ub; SAM80711.1) ^73^; *Ustilago trichophora* (Ut; SPO23210.1) ^74^; *Moesziomyces antarcticus* (Man; GAC76664.1) ^75^; *Moesziomyces aphidis* (Map; ETS62923.1) ^76^. The conserved residues are highlighted in black. The N71 glycosylated site of UmNge1 is marked. (**B**) Phylogenetic analysis of Nge1 orthologs in smut fungi. Protein sequences aligned using Clustal Omega were curated with trimAl (https://ngphylogeny.fr/workflows/alacarte). The phylogenetic analysis was conducted in PhyML 3.0 using the WAG model with 1000 bootstraps. Bootstrap values >50% are indicated at the nodes in 0.1x values for clarity. Clade I: members with the conserved N-glycosylated site are highlighted in blue; Clade II, III: lack N-glycosylation site, Clade III: outgroup. (**C, D**). Nge1 orthologs complement the virulence of Δ*nge1.* Δ*nge1* was complemented by a copy of indicated orthologous alleles SrNge1-HA, UhNge1-HA or UhNge1(S69N)-HA under the control of *UmNGE1* native promoter. The virulence phenotypes of the indicated strains were evaluated. Disease symptoms were assessed as described in the legend of Figure 1C. (**E**) AlphaFold3-generated structures of non-glycosylated and glycosylated Nge1, as well as Nge1-PMEI45 complexes. The non-glycosylated (brown) and glycosylated Nge1 (green) structures were superimposed using ChimeraX (upper panel). The lower panel illustrates the interactions of PMEI45 with both glycosylated and non-glycosylated Nge1. The C-termini of the Nge1 structures are indicated, with N-glycans shown in blue. (**F**) Co-evolutionary escape mechanism between Nge1 and PMEI45, illustrated through interaction dynamics. During the co-evolutionary arms race, maize PMEI45 may have evolved from the PRO-domains or PMEI46 to evade recognition by smut fungal non-glycosylated Nge1, while retaining its conserved function in inhibiting PME19 and PME20 activity. In response, Nge1 has evolved to acquire N-glycosylation as an adaptive modification to overcome the defense mechanism of PMEI45. This modification preserves its interaction with PMEI46 while enabling it to target evasive PMEI45, ultimately releasing PME19 and PME20 to regulate pectin methylesterification levels and presumably loosen the maize cell wall.

We further conducted a phylogenetic analysis using the protein sequences of orthologs to better understand their evolutionary relationship and functional conservation. Nge1 orthologs were categorized into three clades (Figure 6B). Clade I mostly consisted of species with the conserved N-glycosylation site, whereas clade II is characterized by species that lack this site. This phylogeny analysis infers that Nge1 diverged into two distinct groups (clade I and II) that have expanded within smut fungal pathogens. These clades are distantly related to and distinct from the orthologs of clade III (the biocontrol group) ^35^.

Since NYT glycosylation site is mostly conserved within the clade I, this prompted us to investigate whether N-glycosylation-dependent virulence is well-conserved among these orthologs. We assessed the virulence of selected Nge1 orthologs from clades I and II by expressing them in the Δ*nge1* strain, driven by the *UmNGE1* native promoter. The presumably N-glycosylated SrzNge1 ortholog fully complemented the Δ*nge1* virulence function, whereas the UhNge1 ortholog which lacks the N-glycosylation site did not, despite actively expressing at 3 dpi (Figure 6C-D and Figure S6A). To ascertain the importance of N-glycosylation site in UhNge1 virulence, we created an N-glycosylation site by substituting the serine at position 69 with asparagine (S69N). We confirmed that UhNge1(S69N) was N-glycosylated when expressed in *U. maydis*, as evidenced by the mobility shift of the protein upon digestion with PNGase F (Figure S6B). Consequently, this UhNge1(S69N) variant functionally complemented the Δ*nge1* virulence function (Figure 6D).

To compare structural changes between the glycosylated and non-glycosylated Nge1, we superimposed their AlphaFold3-generated structures and detected notable structural changes, particularly at the C-terminus. The N-glycosylation in Nge1 likely induces a conformational shift in the C-terminal intrinsic disorder region, potentially influence its interaction with PMEI45 (Figure 6E). These comprehensive findings suggest that Nge1 exhibits a conserved virulence function that is dependent on N-glycosylation in smut fungi. Our finding further supports the idea that *Um*Nge1 has evolved to acquire glycan modifications to specifically target the evasive PMEI45 as well as PMEI46 to enhance *U. maydis* virulence.

## Discussion

Plants and their associated pathogens are engaged in a continuous coevolutionary arms race, each developing sophisticated strategies to outmaneuver the other. This dynamic interplay drives the evolution of diverse plant defense proteins and modified virulence factors in pathogens, constantly refining their interactions ^36^. Plant cuticles and cell walls act as physical barriers to block pathogen invasion, on which the adapted pathogens must breach or modulate to gain entry. Deletion of the *U. maydis* PME genes does not affect fungal virulence ^37^, but the fungus is capable of increasing PME activity in susceptible maize lines^14^, suggesting the presence of an unexplored invasive strategy. In this study, we uncover a conserved counteradaptive strategy among smut fungi to support fungal infection. Rather than degrading host cell walls and triggering immunity, biotrophic smut fungi employ a novel effector protein Nge1 to hijack PMEIs, alleviating specific PMEs from inhibition to modulate the pectin methylesterification level. Consequently, this changes the dynamics of cell walls and establishes a compatible interaction with plant hosts (Figure 6F).

PMEIs may have originally derived from PRO-domains through whole genome duplication events under selection pressure ^15^. This neofunctionalization of PRO-domains may allow them to avoid being targeted by pathogen virulence factors. This is supported by the observation that although PMEI45 and PMEI46 share high structural similarities and a conserved function in regulating the PME19 and PME20 activities, PMEI45 is able to avoid targeting by non-glycosylated Nge1 orthologs. This finding opens up new avenues for exploring how PMEI45 adapts to evade recognition by non-glycosylated Nge1 while maintaining its conserved function. Resolving the PMEI45 structures by crystallization study could provide insights into the molecular mechanisms driving this evolutionary adaptation, revealing key residues responsible for PMEI45’s evasive mechanisms.

Nevertheless, glycans confer upon effector proteins the ability to acquire new structures and enable novel functions ^38,39^. This molecular adaptation is particularly evident in *U. maydis*, where Nge1 effector proteins are decorated with glycans to fine-tune their interactions in the ongoing co-evolutionary arms race with the host. The dependency on N-glycosylation is specifically observed in Nge1 interaction with PMEI45 but not with PMEI46. Additionally, the ability of UhNge1 (S69N) to restore the virulence phenotype of the Δ*nge1* mutant suggests that glycans confer an adaptive advantage, allowing the glycosylated Nge1 orthologs to overcome PMEI45 evasion (Figure 6F). By exploiting this counteradaptive strategy, smut fungi enhance their ability to successfully colonize the host, underscoring the evolutionary significance of glycan modifications in effector-driven host-pathogen interactions.

The N-glycosylated Cmu1 carries an oligomannose-type N-glycan similar to that of Nge1 but does not interact with PMEI45, indicating that PMEI45’s interaction with Nge1 is not mediated through glycan binding. Instead, a glycan-induced conformational change at Nge1’s C-terminus, which alters the binding pattern, appears to be the driving force. This finding reinforces the idea that N-glycosylation plays a crucial role in shaping protein structure ^40^. Further molecular details on their interaction by crystallization is necessary to elucidate the involvement of specific amino acids. According to phylogenetic analysis of Nge1 orthologs, it is tempting to speculate that during the coevolutionary arms race, these orthologs undergo selection pressure, evolving either at the post-translational level via glycan modification (Clade I) or at the DNA level (Clade II) to counteract newly evolved PMEIs in their specific hosts. This may enable them to achieve a similar function of impacting host cell wall dynamics and favor colonization.

Several studies have reported contradictory subcellular locations for the processing of PRO-PMEs. One study reported that the PRO regions retain unprocessed in the Golgi until cleavage is completed before exporting PMEs ^41^. In contrast, others argue that the processing of PRO-PMEs could occur at the cell wall ^42–44^. To date, there is no direct evidence that supports PRO-PME processing in the apoplast. Such uncertainty raises the possibility that PRO19 or PRO20 might also be the bona fide targets of Nge1 in the apoplast. However, based on the N-glycosylation-dependent virulence function of Nge1, we propose a role for Nge1 in interfering with the action of PMEI45 on PME activities.

Our finding cracks the mystery of how fungi outfox the defensive strategies of multiple PMEIs in the host, all by leveraging just one Nge1 effector with the glycan decoration. Moving forward, uncovering the specific amino acids involved in these interactions and elucidating the precise mechanisms by which N-glycosylation facilitates this process will undoubtedly deepen our understanding and pave the way for further discoveries. Enhancing the degree of highly methylesterified pectin through the overexpression of *PMEI* genes could potentially strengthen maize resistance against biotrophic pathogens, presenting promising avenues for disease control strategies.

## Materials and Methods

### Plant Materials and Growth

Seven-day old maize seedlings Honey 236, a Taiwan cultivar, used for *U. maydis* infections were grown under controlled conditions (15 h/9 h light/dark cycle, 28/25 °C). 3-week old *Nicotiana benthamiana* seedlings grown under a 14 h/10 h light/dark cycle, 25/22 °C, were used for *Agrobacterium* infiltrations.

### Microbial Strains and Growth Conditions

The haploid solopathogenic *U. maydis* SG200 ^26^, was grown at 28°C on potato dextrose agar plates (PDA; 2.4% potato dextrose broth and 2% agar) and in YEPSL liquid medium (0.4% yeast extract, 0.4% peptone, and 2% sucrose). Hygromycin (400 μg/ml) and carboxin (4 μg/ml) were filter sterilized and supplemented to PDA for positive selection. *Saccharomyces cerevisiae* AH109 was grown at 28°C in yeast extract peptone dextrose medium (YPD; 1% yeast extract, 2% peptone, 2% agar, and 2% glucose). *Agrobacterium* strain GV3101 was grown at 28°C in DYT medium (1.6% tryptone, 1% yeast extract, and 0.5% NaCl) supplemented with rifampicin (40 μg/ml) and gentamycin (50 μg/ml).

### Plasmids and Strains

*U. maydis* strains generated in this study are documented in Table S1. Plasmids constructed using standard cloning methods ^45^ or Gibson Assembly ^46^ are listed and described in Table S2. Primers for each plasmid generation are provided in Table S3. For gene integration into the *ip* locus, plasmids containing upstream and downstream regions, and a carboxin resistant *ip* allele (*ip*R) ^47^ were linearized and subsequently inserted into the genome via homologous recombination-based gene integration. The *U. maydis* transformation and genomic DNA isolation procedures were performed as described in a previous study ^48^. Positive transformants were verified by Southern blot analysis.

### U. maydis Infection

*U. maydis* strains cultured in YEPSL until reaching an OD_600_ of 0.8 were adjusted to a final OD_600_ of 1.0 using MilliQ water before inoculation onto 7-day-old maize seedlings. Disease symptoms were scored based on established criteria at 8 days post-infection (dpi) ^26^. The disease severity index (DI) was calculated ^49^, and significant differences between strains indicated in the figures were determined using a two-tailed unpaired Student’s *t*-test.

### Gene Expression Analysis

qRT-PCR analysis of *NGE1* was conducted as previously described with minor adjustments ^49^. Briefly, total RNAs of SG200 cells (OD_600_ ∼0.8) and SG200-infected maize leaves were extracted using TRIzol (Invitrogen#15596018), DNase treated (TURBO DNA-free™ Kit, Invitrogen#AM1907), and reverse transcribed (SuperScript® III First-Strand Synthesis SuperMix, Invitrogen#18080400). cDNA was subjected to qRT-PCR analysis using primer pairs listed in Table S1C. The expression levels of *U. maydis* peptidyl-prolyl isomerase (PPI) were used for normalization. Relative expression values were calculated using the 2^−ΔΔCt^ method ^50^.

### *In-vitro* Secretion and Deglycosylation

*U. maydis* cells were cultured to an OD_600_ of 0.6 and harvested. Cell pellets were lysed in 1x sample buffer (50 mM Tris-HCl, 2% SDS, 10% glycerol, 100 mM DTT, 0.01% bromophenol blue, pH 6.8), culture supernatants were TCA-precipitated to obtain total and secreted proteins, respectively. Proteins were immunoblotted using mouse antibodies against HA (dilution 1:5000; Yao-Hong Biotech., Taiwan; #YH80007) or tubulin (dilution 1:7500; Sigma, #T6199) and a goat secondary antibody conjugated with horseradish peroxidase (dilution 1:50,000; Yao-Hong Biotech., Taiwan; #AS111772). For deglycosylation, supernatant fractions were treated with PNGase F (New England Biolabs #P0704) in a final volume of 25 μl, following the manufacturer’s protocol.

### *N*-Glycosylation Site Identification

Nge1-mCherry-HA proteins were immunoprecipitated from the concentrated culture supernatant sample using anti-HA magnetic beads (Sigma) and visualized on a silver-stained SDS page. The Nge1-mCherry-HA band was excised, digested in-gel with trypsin, desalted using a C18 cartridge, and then subjected to PNGase-F deglycosylation. Peptides were desalted again, lyophilized, and analyzed by NanoLC-nanoESI-MS/MS on an EASY-nLC™ 1200 system coupled to a Thermo Orbitrap Fusion Lumos mass spectrometer with a Nanospray Flex™ ion source. The MS analysis was conducted over a mass range of m/z 350 to 1600 with a resolution of 120,000 and an ion count target of 2 × 10⁵. Fragmentation was achieved through Higher-Energy Collisional Dissociation (HCD) with a normalized collision energy set to 30 eV. The MS² analysis was performed at a resolution of 30,000, with an ion count target of 5 × 10⁴ and a maximum injection time of 54 ms. Peptide identification was done using Proteome Discoverer software with SEQUEST and Mascot search engines against the *U. maydis* database (UniProt), with deamidation modification set.

### Protein Purification

The culture supernatant of *U. maydis* cells constitutively expressing secreted Nge1-His were collected and buffer-exchanged to binding buffer (25 mM Tris, 0.3 M NaCl, 10 mM Imidazole, pH 7.5). Subsequently, the sample was passed through a 5 ml HisTrap column (Cytiva) connected to an AKTA Pure 25M system (GE). The column was washed with wash buffer (25 mM Tris, 0.3 M NaCl, 20 mM Imidazole, pH 7.5), and proteins were eluted using an elution buffer (25 mM Tris, 0.3 M NaCl, 250 mM Imidazole, pH 7.5) with a linear gradient of imidazole concentration and collected in a fraction collector. Finally, the purified proteins were analyzed using Coomassie blue-stained SDS PAGE.

### Identification of N-glycan Patterns

Nge1-His (5 μg), purified from *U. maydis* culture supernatant, was digested with chymotrypsin, while Cmu1-His (5 μg) was digested with trypsin and Asp-N endopeptidase. The resulting peptides were detected by LC-ESI-MS on an Orbitrap Fusion mass spectrometer (Thermo Fisher Scientific, San Jose, CA) equipped with an EASY-nLC 1200 system and EASY-spray source. Full-scan MS condition: mass range m/z 375-1800 with an automatic gain control (AGC) target of 5 × 10⁵ with lock mass, resolution 60,000 at m/z 200, and maximum injection time of 50 ms. The MS² was run in top speed mode with 3s cycles with Collision-Induced Dissociation (CID) and HCD. Glycan identification was performed using Byonic software 4.2.10 (Protein Metrics) ^51^ with a glycan database containing 132 entries.

### Apoplastic Localization of Nge1-mCherry

*U. maydis*-infected leaf segments, approximately 3 cm in length below the infection sites, were collected at 3 dpi, washed with sterile water, and pat-dried. Plasmolysis was induced by immersing the leaf segments in 0.8 M mannitol and subjecting them to a 5-minute vacuum infiltration (Eppendorf® Vacufuge Plus Complete System). The samples were kept at room temperature until confocal microscopy (Zeiss LSM880). The mCherry signal was visualized with excitation at 561 nm and emission at 570–630 nm.

### Target Identification of Nge1

Culture supernatants from *U. maydis* strains expressing HA-tagged Nge1 and N71Q were concentrated 30-fold using 3 kDa cutoff centrifugal filters (GE) and buffer-exchanged to 25 mM Tris-HCl (pH 7.5). Bait proteins were immobilized onto anti-HA magnetic beads and washed with binding buffer A (25 mM Tris-HCl, 0.3 M NaCl, and 0.1% NP-40, pH 7.5). Δ*nge1*-infected maize leaves at 4 dpi were immersed in calcium buffer (0.2 M CaCl_2_, 5 mM NaAcetate, pH 4.3) and apoplastic fluids were extracted using vacuum-centrifugation method ^52,53^. The apoplastic fluid was buffer-exchanged to buffer B (25 mM Tris-HCl, 0.3 M NaCl, 0.1% NP-40, and 5 mM NaAcetate, pH 7.0) before incubation with the bait-containing beads. After overnight incubation at 4°C, bound proteins were washed with buffer B lacking NaAcetate, and then subjected to silver-stained SDS-PAGE analysis. Distinct bands were excised, trypsin digested, and subjected to LC-MS/MS analysis.

### Yeast Two-hybrid Analysis

Yeast two-hybrid assays were conducted following the protocol described ^54^. Briefly, positive transformants were selected on SD medium lacking leucine (L) and tryptophan (W). To assess protein interactions, yeast transformants were allowed to grow on SD plates lacking LW, histidine (H), and adenine (A), or lacking LWH and supplemented with 3-amino-1,2,4-triazole (3-AT) for 3 to 5 days at 28°C. The expressions of proteins were detected by immunoblotting.

### Bimolecular Fluorescence Complementation (BiFC)

*A. tumefaciens* GV3101 strains carrying desired plasmids were co-infiltrated into 3 to 4-week-old *N. benthamiana* leaves. At 4 dpi, YFP signal with excitation at 514 nm and emission at 541 nm was visualized by confocal microscope. Under the same parameter setting, images of leave cells were acquired, and total intensities were quantified and normalized with the cell wall autofluorescence observed at excitation 405 nm and emission at 427 nm. Significant differences in YFP signals between the two groups were determined using a two-tailed unpaired Student’s *t*-test. Split-YFP fusion proteins were immunoblotted using rabbit polyclonal antibodies against GFP (dilution 1:5000; Yao-Hong Biotech., Taiwan; #YH80006) and a goat secondary antibody conjugated with horseradish peroxidase (dilution 1:25,000; Yao-Hong Biotech., Taiwan; #AS09602).

### FoMV-induced Gene Overexpression

Constitutive gene expression in maize was performed using the Foxtail mosaic virus pV101 vector-based system with slight modifications ^33^. *Agrobacterium* carrying pV101 constructs expressing desired proteins of interest were infiltrated into *N. benthamiana* leaves. Viral sap was collected at 10 dpi by grinding leaf tissue in 1x PBS, pH 7.4. This sap was rub-inoculated on 6-day-old maize seedlings pre-incubated in darkness for a day, using 600-mesh carborundum powder. Two rounds of viral sap multiplication were conducted, and symptomatic maize leaves were harvested at 10 dpi to prepare 2 μm thick cross-sections. Maize seedlings inoculated with viral sap three days prior were used for *U. maydis* infection.

### Immunostaining of Pectins

Immunostaining of pectins in maize leaf cross-sections followed a standardized protocol^55^. Cross section slides were treated with 0.02% calcofluor white for 10 minutes, washed with water and 1x TBS-T, and then blocked with 2% (w/v) non-fat skim milk in 1x TBS-T for 75 minutes. Pectin-specific rat monoclonal antibodies LM19 and JIM7 (I-ELD001 and I-ELD005 from Kerafast, Inc., Boston, USA) and goat anti-rat secondary antibody conjugated to Alexa-Fluor 488 (dilution 1:50, ab150157) were used for immunostaining.

### Bioinformatic Analyses

The Nge1 amino acid sequence was retrieved from the fungal genome resource (https://mycocosm.jgi.doe.gov/Ustma2_2/Ustma2_2.home.html) ^26^. Domain prediction was conducted using InterPro (https://www.ebi.ac.uk/interpro/) ^25^. AlphaFold structures were obtained from UniProt (https://www.uniprot.org/) ^56,57^ and superimposed using ChimeraX (https://www.cgl.ucsf.edu/chimerax/download.html) ^58^. TM align was used to compare the structures of two proteins (https://zhanggroup.org/TM-align/) ^31^. The AlphaFold structures of PMEI45-Nge1 complex were generated using AlphaFold 3 (https://alphafoldserver.com) ^59^. SignalP 6.0 (https://dtu.biolib.com/SignalP-6) was utilized for signal peptide prediction ^60^. ApoplastP (https://apoplastp.csiro.au/) was employed to predict the apoplast localization of Nge1 ^27^. N-glycosylation sites were predicted using the NetNglyc server (http://www.cbs.dtu.dk/services/NetNGlyc/) ^61^. Nge1 orthologs were identified via Blastp analysis on NCBI against the non-redundant database (https://blast.ncbi.nlm.nih.gov/Blast.cgi). Sequence alignments were generated using Clustal Omega (https://www.ebi.ac.uk/jdispatcher/msa/clustalo) ^62^ and visualized with GeneDoc (https://genedoc.software.informer.com/2.7/). Phylogenetic analysis was conducted using PhyML 3.0 (http://www.atgc-montpellier.fr/phyml/) ^63^ with the Smart Model Selection method ^64^, and the resulting tree was visualized and annotated using iTOL (https://itol.embl.de/) ^65^.

## Acknowledgments

We are grateful to Kostya Kanyuka (The National Institute of Agricultural Botany [NIAB], Cambridge, U.K.) for providing the FoMV PV101 vectors. We acknowledge Drs. Erh-Min Lai, Chih-Hang Wu, and Li-Hung Chen for their valuable suggestions and comments. We thank Dr. Erh-Min Lai for kindly sharing plasmids for generating BiFC constructs. We appreciate technical assistance and services provided by the staff of Academia Sinica Glycoscience Core Facility (funded by AS-CFII-112-102 project) and the core laboratories IPMB, including Proteomics, DNA Sequencing, Live-Cell-Imaging, and Cell-Biology EM. This work is supported by NSTC grant (MOST 111-2311-B-001-026-MY3) and the TIGP Ph.D. fellowship.

## Author Contributions

LSM and CB conceived, designed, and wrote the manuscript with input from all authors. CB performed most of the experiments. MQC conducted Y2H experiments and assisted with pectin-staining. WLT initiated the VOX assay. OKT provided technical support, advised on pectin methylation analysis, and critically discussed the manuscript. All authors contributed to the discussion of the results and reviewed the manuscript.

---

**Table S1**. *U. maydis* strains used in this study

**Table S2**. Plasmid constructs used in this study

**Table S3**. Primers used for plasmid constructs in this study

**Table S4**. The accessions of maize type-I PMEs and PMEI proteins

**Table S5**. Y2H construct information

**Figure S1.**
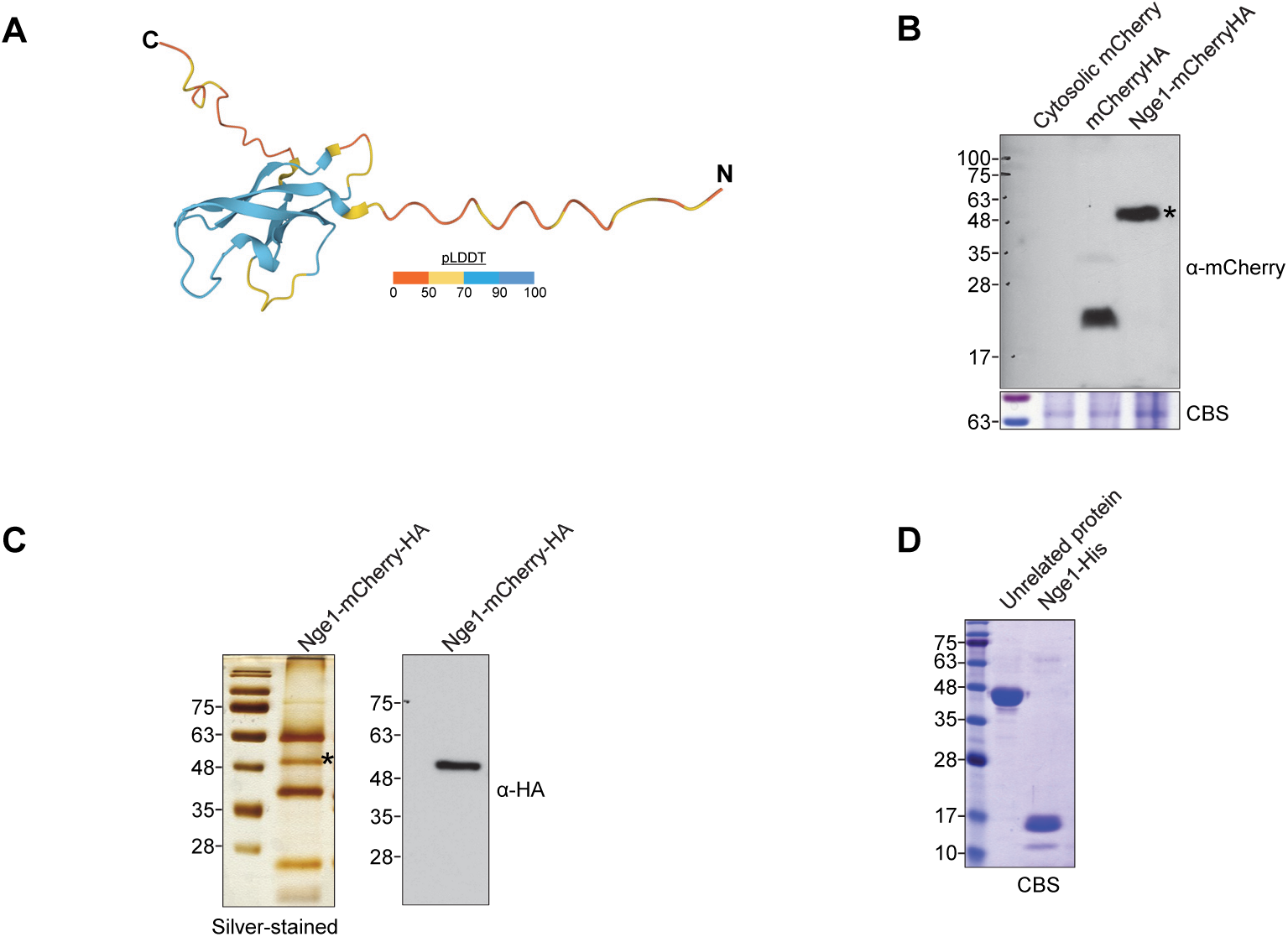
Protein detection and analysis. (**A**) The AlphaFold2 predicted structure of Nge1 is shown with pLDDT indicating local confidence levels. (**B**) Immunoblot detection of Nge1-mCherry in the apoplastic fluid. TCA-precipitated-apoplastic fluid extracted from maize leaves infected with the strains SG200Δ*nge1_*Nge1-mCherryHA, SG200_mCherry (producing non-secreted mCherry) or SG200_SP_cmu1_-mCherryHA was analyzed using immunoblotting. Cytosolic mCherry served as a negative control, and SP-mCherryHA served as a positive control. Coomassie blue staining (CBS)-membranes were used to verify loading. Asterisks indicate the full length of fusion Nge1-mCherryHA (⁓51 kDa). (**C**) Immunoprecipitation of Nge1-mCherryHA. Nge1-mCherryHA fusion proteins were immunoprecipitated from the culture supernatant of SG200 strain constitutively expressing Nge1-mCherryHA using anti-HA magnetic beads. The proteins were visualized on a silver-stained SDS-PAGE and immunoblotted against HA-epitopes. Asterisk indicate the Nge1-mChery-HA protein band. (**D**) Affinity purification of Nge1-His. Nge1-His proteins were purified from the *U. maydis* culture supernatant of SG200Δ*nge1_*Nge1-His using a HisTrap column. The purified proteins were then separated by SDS-PAGE and visualized with CBS.

**Figure S2.**
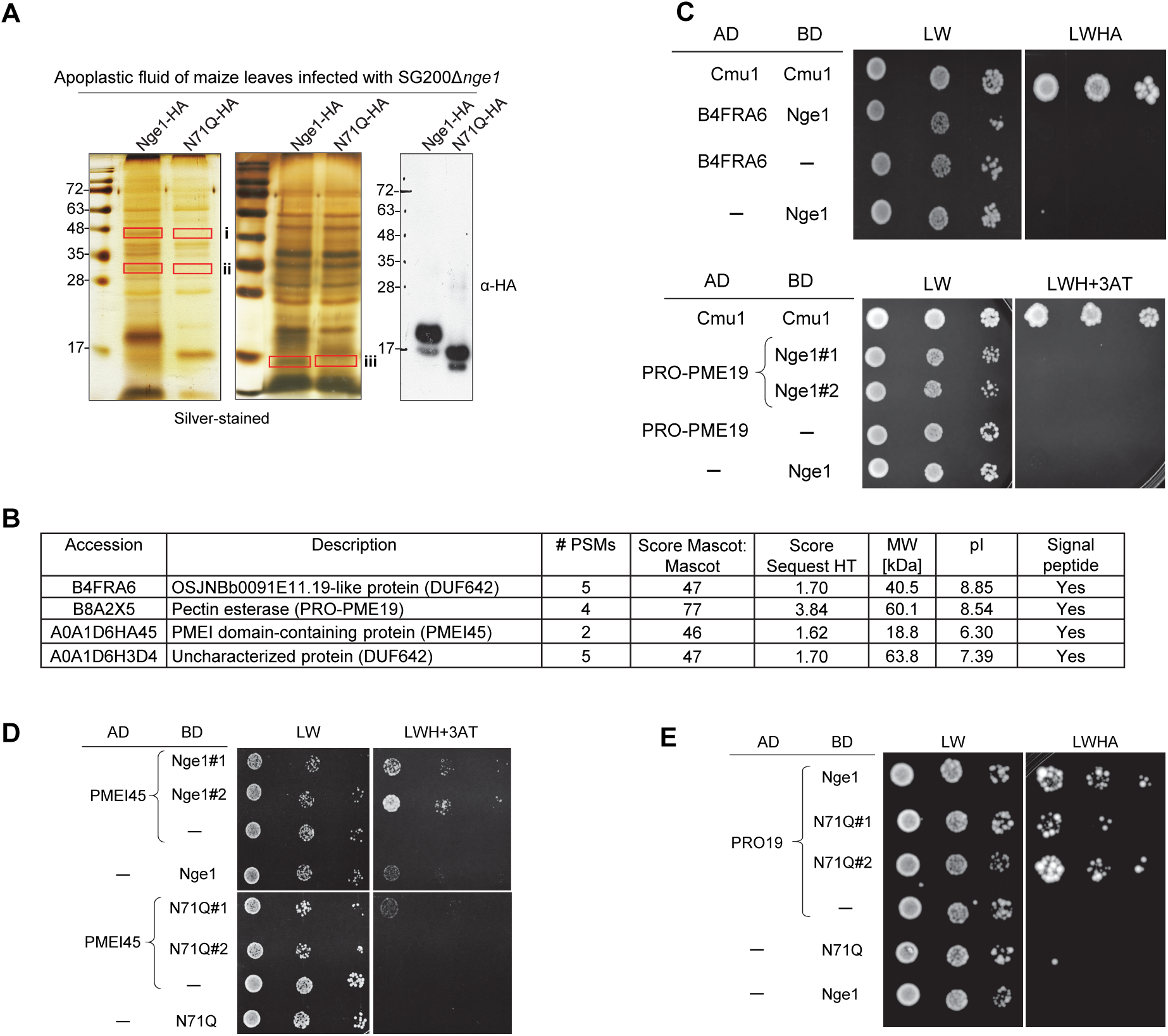
Identification of maize apoplastic proteins pull down by Nge1. (**A**) Apoplastic fluid from G200Δ*nge1*-infected maize leaves was incubated with Nge1-HA or N71Q-HA immobilized on anti-A magnetic beads. The bound proteins were eluted, separated and visualized by silver-stained SDS-AGE. The protein bands i, ii, and iii, marked by a red rectangles, were excised and subjected to LC-MS/MS analysis. (**B**) Detailed information of pectin methylesterase (PME) activity-related candidates identified in the LC-MS/MS analysis of Nge1-interacting proteins are shown. PSM: peptide-spectrum match. (**C-E**) Yeast-two hybrid analysis for the interaction of Nge1 with the potential candidates. (**C**) DUF642 and full-length PME19 are not the targets of Nge1. (**D**) Nge1 targets PMEI45. (**E**) Nge1 nteracts with PRO-domain of PME19. Yeast transformants containing two plasmids expressing ndicated proteins fused to GAL4 activation domain (AD) or binding domain (BD) without signal peptide ere grown on SD-Leu/-Trp (LW), SD-Leu/-Trp/-His/-Ade (LWHA), or SD-LWH containing 3mM of 3AT 3-amino-1,2,4-triazole) plates for 2-3 days. ̶, empty vector; Chorismate mutase (Cmu1) served as the ositive control; Similar results were observed in at least two independent experiments. Detailed nformation on the protein regions used in the Y2H analysis is available in Table S5.

**Figure S3.**
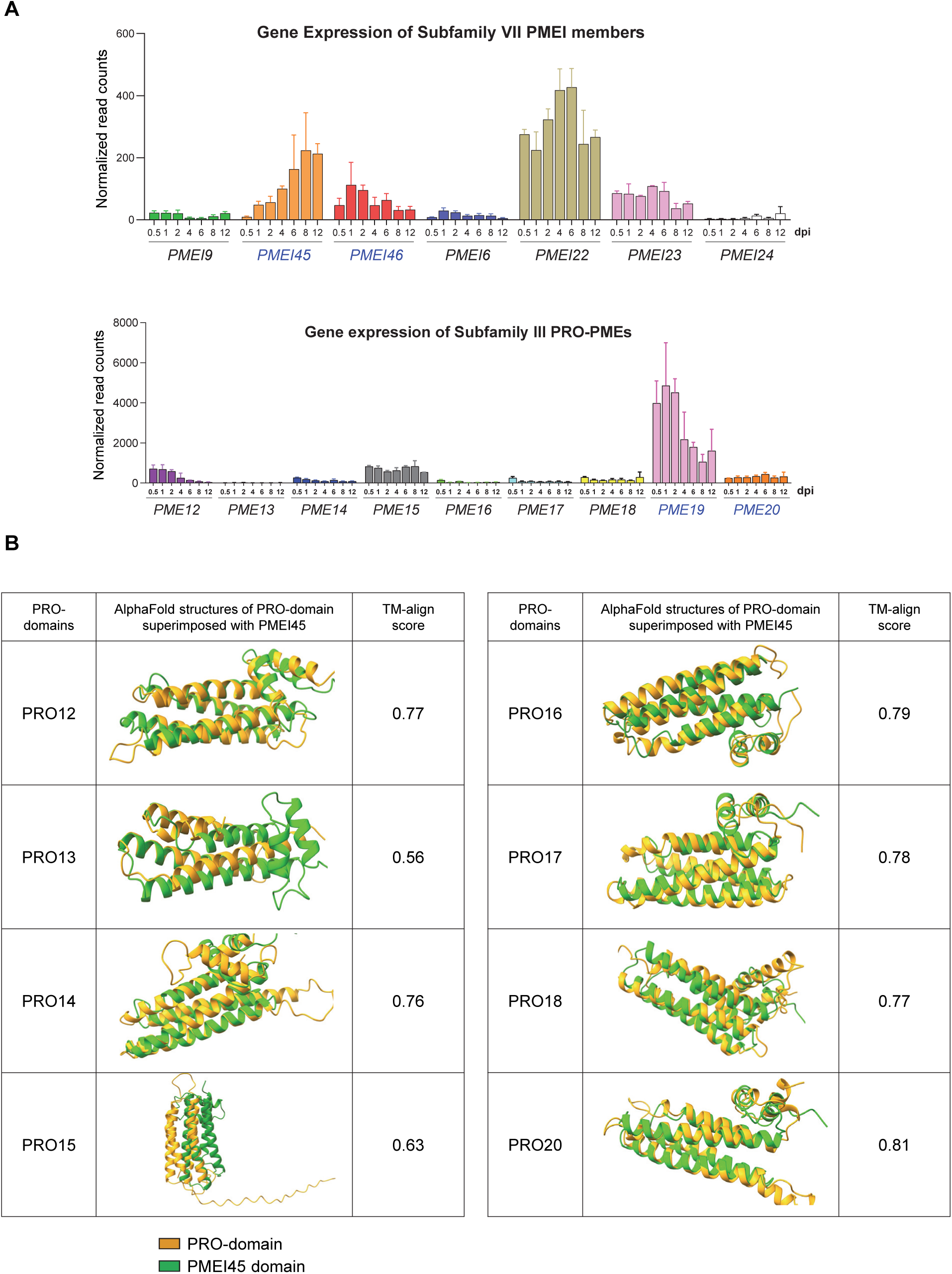
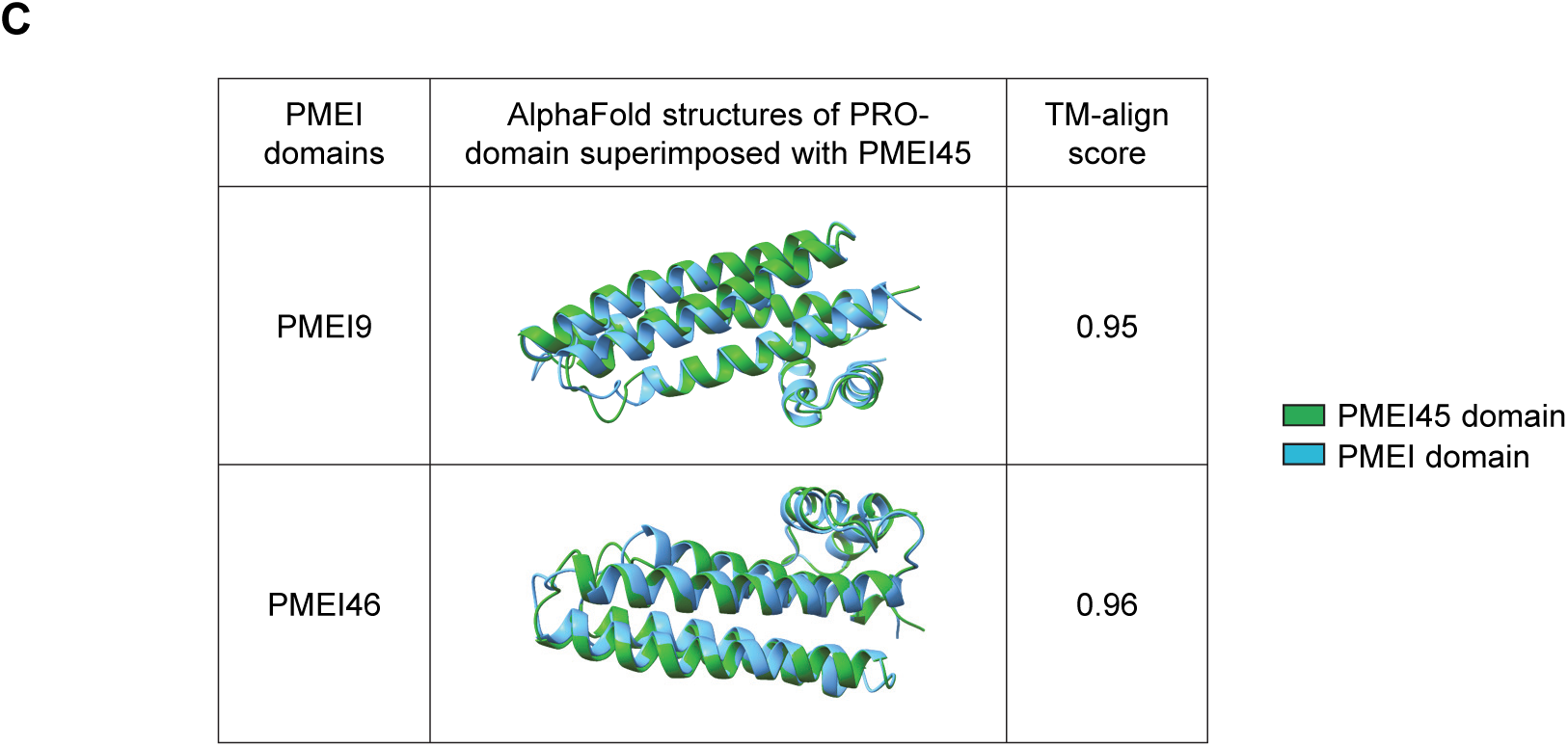
RNAseq expression profile of subfamilies III and VII members during *U. maydis* infection and their AlphaFold structure aligns with PMEI45. (**A**) RNAseq expression profiles of subfamily III and VII members. Data were compiled from a public dataset generated by Lanver et. al. 2018 (https://doi.org/10.1105/tpc.17.00764). Dpi: days post infection. Superimposition of indicated PRO-domains from PMEs (**B**) or PMEIs (**C**) with the PMEI45 domain. The AlphaFold structures were generated using AlphaFold2 and superimposed using ChimeraX. Structural similarity was analyzed by TM-align, with TM-scores shown.

**Figure S4.**
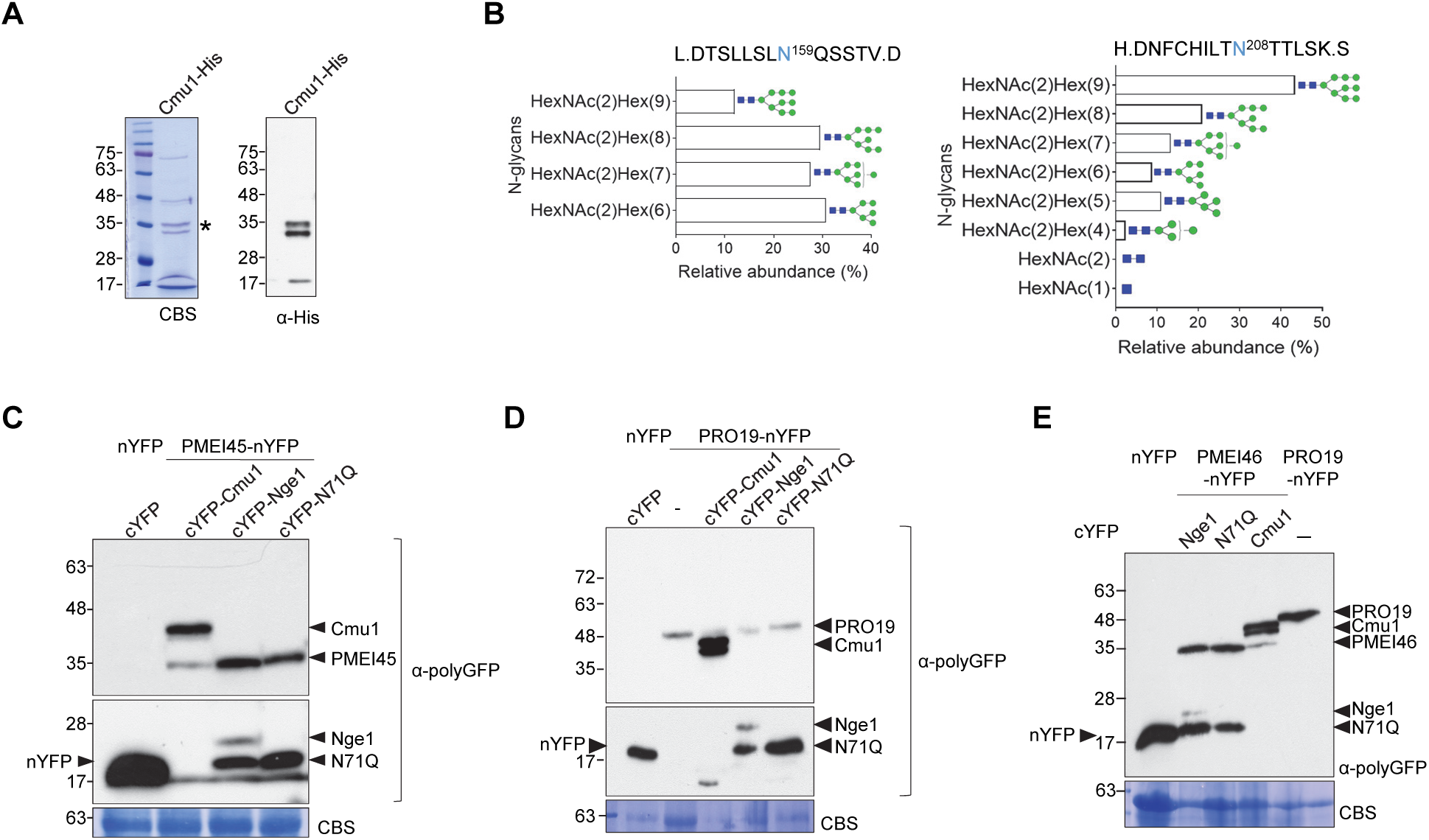
(**A-B**) ***U. maydis* Cmu1 effector is decorated with oligomannose-type N-glycans. (A**) Affinity purification of Cmu1-His. Cmu1-His proteins from the *U. maydis* culture supernatant were separated by SDS-PAGE and visualized with CBS and immunoblotting. Asterisks indicate the Cmu1-His protein (> expected 30.8kDa). (**B**) Detection of N-glycans at N159 and N208 in Cmu1 effector by LC-MS/MS. The graphs illustrates the relative abundance percentage of detected oligomannose type N-glycan forms on two glycopeptides, LDTSLLSLNQSSTVD; HDNFCHILTNTTLSKS in a replicate. HexNAc and Hex is proposed as GlcNAc and mannose respectively. Blue square, N-acetylglucosamine; Green circle, mannose. The number of sugar molecules is indicated. (**C-E**) **Protein expression in tobacco plants for BiFC assays.** Immunoblot detection of PMEI45 (**C**) or PRO19 (**D**) or PMEI46 (**E**) fused to nYFP at the C-terminus, co-expressed with Cmu1, Nge1, or N71Q fused to cYFP at the N-terminus in tobacco plants. TCA-precipitated proteins extracted from tobacco leaves infiltrated with Agrobacterium expressing the indicated proteins were analyzed by immunoblotting using polyGFP antibodies. Co-expression of nYFP and cYFP split or PRO19-nYFP alone served as negative controls. Molecular weight of mature proteins, nYFP: ⁓19.7 kDa: cYFP: ⁓7.3 kDa; PMEI45-nYFP: ⁓36.1 kDa; PRO19-nYFP: ⁓38.5 kDa (one putative glycosylation site); PMEI46-nYFP: ⁓36.2 kDa, cYFP-Cmu1: ⁓37.2 kDa; cYFP-Nge1: >18 kDa (N-glycosylated); cYFP-N71Q: ⁓18 kDa. The protein bands for the respective proteins are labelled. CBS of membranes was used to verify protein loading.

**Figure S5.**
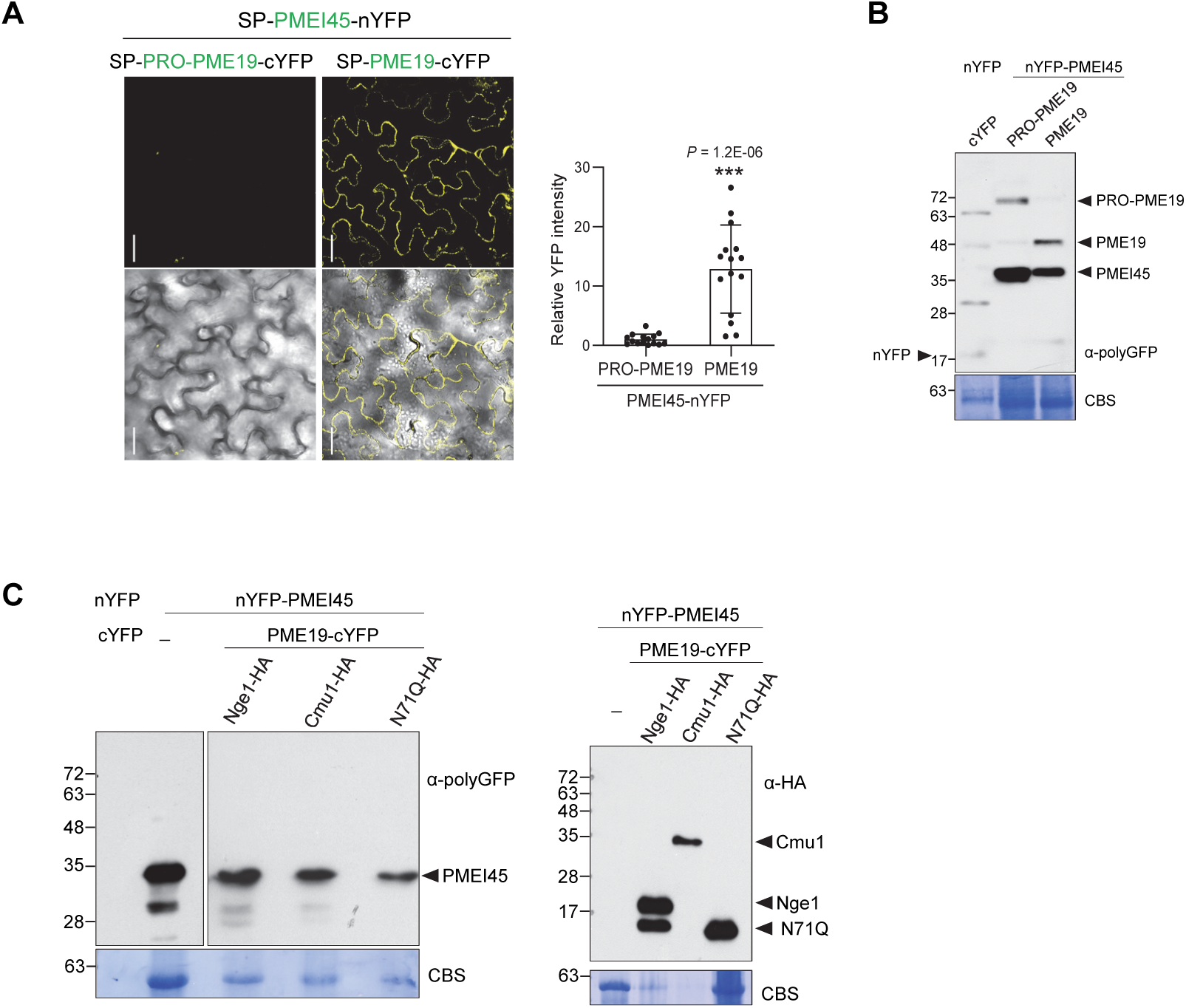
BiFC analysis of the PMEI45-PME19 interaction and protein expression in tobacco plants. (**A**) PMEI45 interacts with PME19 enzymatic domain. BiFC analysis of PMEI45 interaction with PME19 or PRO-PME19 *in planta*. The graph depicted the fold differences relative to the control expressing PRO-PME19. Values indicate mean ± SD from total samples obtained from three independent biological assays and dots depict the values of individual samples across these replicates. Significant differences compared to PRO-PME19 as determined by a two-tailed unpaired Student’s *t*-test. All bars 50 μm. (**B-C**) Immunoblot detection of indicated proteins expressed in tobacco plants. TCA-precipitated proteins extracted from tobacco leaves infiltrated with Agrobacterium expressing the indicated proteins were analyzed by immunoblotting using polyGFP antibodies. Molecular weight of mature proteins, nYFP: ⁓19.7 kDa: cYFP: ⁓7.3 kDa; PMEI45-nYFP fusion: ⁓36.1 kDa; The protein bands for the respective proteins are labelled. CBS of membranes was used to verify protein loading. (**B**) Immunoblot detection of PMEI45 fused to nYFP at the N-terminus, co-expressed with PRO-PME19 or PME19 fused to cYFP at the C-terminus in tobacco plants. PRO-PME19-cYFP: >64.9 kDa (3 putative glycosylation sites); PME19-cYFP: >44 kDa (2 putative N-glycosylation sites). (**C**) Detection of HA-tagged Nge1 or Cmu1 or N71Q co-expressed with nYFP-PMEI45 and PME19-cYFP in tobacco plants by immunoblotting using HA antibodies and polyGFP antibodies.

**Figure S6.**
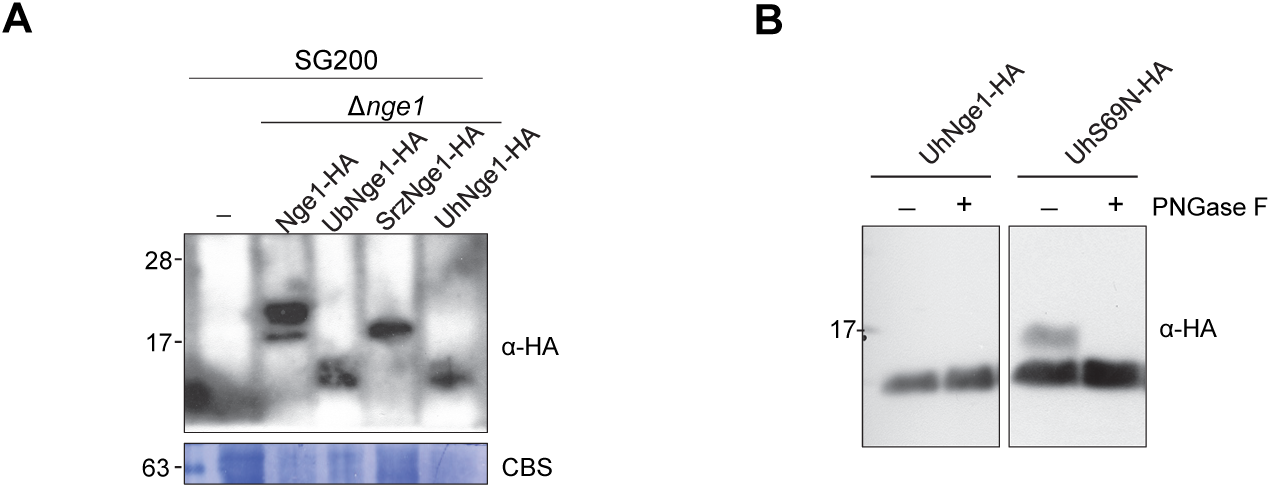
Nge1 orthologs complemented the virulence of *U. maydis* Δ*nge1*. (**A**) Detection of Nge1 ortholog proteins *in planta* at 3dpi. TCA-precipitated proteins from maize leaves infected with the indicated strains were subjected to immunoblotting using anti-HA antibodies. UbNge1: ⁓ 8.9 kDa; SrzNge1: ⁓9.7 kDa; UhNge1: ⁓ 8.9 kDa. Coomassie-stained membranes (CBS) served as a loading control. (**B**) UhNge1(S69N) is glycosylated. Deglycosylation of UhS69N. The supernatant fractions of *U. maydis* constitutively expressing UhS69N-HA were treated with or without PNGase F and subjected to immunoblotting. UhNge1 served as a negative control.

